# Network Proteomics of the Lewy Body Dementia Brain Reveals Presynaptic Signatures Distinct from Alzheimer’s Disease

**DOI:** 10.1101/2024.01.23.576728

**Authors:** Anantharaman Shantaraman, Eric B. Dammer, Obiadada Ugochukwu, Duc M. Duong, Luming Yin, E. Kathleen Carter, Marla Gearing, Alice Chen-Plotkin, Edward B. Lee, John Q. Trojanowski, David A. Bennett, James J. Lah, Allan I. Levey, Nicholas T. Seyfried, Lenora Higginbotham

## Abstract

Lewy body dementia (LBD), a class of disorders comprising Parkinson’s disease dementia (PDD) and dementia with Lewy bodies (DLB), features substantial clinical and pathological overlap with Alzheimer’s disease (AD). The identification of biomarkers unique to LBD pathophysiology could meaningfully advance its diagnosis, monitoring, and treatment. Using quantitative mass spectrometry (MS), we measured over 9,000 proteins across 138 dorsolateral prefrontal cortex (DLPFC) tissues from a University of Pennsylvania autopsy collection comprising control, Parkinson’s disease (PD), PDD, and DLB diagnoses. We then analyzed co-expression network protein alterations in those with LBD, validated these disease signatures in two independent LBD datasets, and compared these findings to those observed in network analyses of AD cases. The LBD network revealed numerous groups or “modules” of co-expressed proteins significantly altered in PDD and DLB, representing synaptic, metabolic, and inflammatory pathophysiology. A comparison of validated LBD signatures to those of AD identified distinct differences between the two diseases. Notably, synuclein-associated presynaptic modules were elevated in LBD but decreased in AD relative to controls. We also found that glial-associated matrisome signatures consistently elevated in AD were more variably altered in LBD, ultimately stratifying those LBD cases with low versus high burdens of concurrent beta-amyloid deposition. In conclusion, unbiased network proteomic analysis revealed diverse pathophysiological changes in the LBD frontal cortex distinct from alterations in AD. These results highlight the LBD brain network proteome as a promising source of biomarkers that could enhance clinical recognition and management.

## Introduction

Lewy body dementia (LBD), a class of disorders comprising Parkinson’s disease dementia (PDD) and dementia with Lewy bodies (DLB), is the second most common cause of dementia worldwide and is currently without cure or effective mitigating therapies [1]. The identification of reliable biomarkers of its aggressive cognitive and neuropsychiatric symptoms is critical to advancing the clinical management of LBD. Yet, LBD biomarker discovery has proven challenging, in large part due to the complexity and often overlapping pathophysiology driving its dementia, psychosis, and mood disturbances. Pathological evidence has linked the cognitive and neuropsychiatric manifestations of LBD to the diffuse deposition of α-synuclein-rich Lewy bodies (LBs) throughout the limbic and neocortex [2-6]. However, a growing number of studies across multiple disciplines, including genetic, clinicopathological, imaging, and biofluid analyses, suggests LBD pathophysiology encompasses a diverse array of corticolimbic processes extending beyond synuclein accumulation and neuronal loss. Among these implicated mechanisms are mitochondrial dysfunction, neuroinflammation, aberrant cholinergic and other neurotransmitter activity, and synaptic dysregulation [1, 7-9]. This increasingly complex pathophysiological landscape indicates multiple molecular signatures may be necessary for effective LBD biomarker development.

The genetic and molecular overlap LBD shares with Alzheimer’s disease (AD) and other neurodegenerative disorders further complicates the ability to unravel its key pathophysiological signatures. Genome-wide association studies (GWAS) have identified a significant amount of polygenic overlap between LBD and AD [7]. Meanwhile, according to certain clinicopathological estimates, over 50% of those with LBD feature concurrent accumulation of the extracellular amyloid-beta (Aβ) plaques and tau neurofibrillary tangles (NFTs) that comprise the core of AD neuropathology [10]. Furthermore, limbic-predominant TAR DNA-binding protein 43 (TDP-43) and other pathological inclusions are frequently detected in both LBD and AD [11]. This overlapping pathology has a marked impact on diagnostic accuracy, clinical trial stratification, disease prognosis, and therapeutic development [10-13]. Thus, research strategies designed to better define not only unique signatures of LBD pathophysiology but also its degree of overlap with other neurodegenerative diseases are critical to the discovery of biomarkers that meaningfully advance its diagnosis, monitoring, and treatment.

Network-based proteomics, which quantifies global pathophysiological changes in complex biological samples [14], is a tool designed to address many of these challenges. This data-driven approach organizes large proteomic datasets into groups or “modules” of proteins with similar expression patterns across individual samples. These co-expression modules are often enriched with markers specific to certain cell types, molecular functions, and organelles, providing insights into the diverse pathophysiological alterations reflected in the disease specimen. We have previously used this approach to define and characterize the network of complex protein pathophysiology within the brain tissues of those with pathologically defined AD [15-22]. These AD-associated modules and their hub proteins have proven highly reproducible across different tissue cohorts and brain regions, allowing us to generate a large consensus AD brain network across hundreds of corticolimbic tissues [17]. In sum, our consensus findings have 1) enhanced understanding of neuronal and non-neuronal pathophysiology in the AD brain; 2) provided a strong molecular framework for network-level comparison to other neurodegenerative diseases, and 3) served as a strong foundation for panel-based biomarker discovery in cerebrospinal fluid and plasma [23, 24]. Furthermore, these proteomic networks have revealed significant disease-related alterations not reflected at the transcriptomic level [15-17, 20].

In the current study, we apply an unbiased co-expression network proteomic approach to the study of corticolimbic alterations in the brains of those with pathologically defined LBD, establishing a global systems-based framework of the protein-level changes underlying neurodegeneration in these tissues. In addition, we perform a network-level comparison of the LBD and AD brain proteomes. Our results reveal protein co-expression alterations throughout a diverse range of pathophysiological systems in the LBD brain, including presynaptic signatures distinct from those observed in the AD proteomic network. We also demonstrate how α-synuclein (SNCA) plays a critical “bottleneck” role in the network-level communication among these synaptic signatures. Finally, we underscore the utility of proteomic network analysis in examining not only divergent changes but also overlapping features of LBD and AD, highlighting signatures capable of stratifying LBD cases with low versus high burdens of amyloid co-pathology. Overall, this approach offers a systems-based foundation for the discovery of protein biomarkers that reflect the unique and complex pathophysiology of LBD.

## Results

### Differential expression analysis demonstrates robust protein alterations in the LBD brain

The main objective of this study was to perform unbiased co-expression network analysis of LBD brain tissues to better define global pathophysiological alterations in cortical regions and compare these findings to the AD brain network proteome. All brain tissues were derived from a pathologically well-characterized autopsy collection within the University of Pennsylvania (UPenn) Alzheimer’s Disease Research Center (ADRC). Bulk tissue homogenates from the dorsolateral prefrontal cortex (DLPFC) of cases with neuropathologically confirmed diagnoses of healthy control (*n*=47), Parkinson’s disease (PD; *n*=33), PDD (*n*=47), and DLB (*n*=11) were included in our initial network analysis. The frontal cortex was examined as it is often affected in the diffuse corticolimbic LB accumulation found in LBD and is routinely scored in its neuropathological diagnosis [25]. In addition, frontal executive deficits are commonly among the first symptoms observed in LBD [26], suggesting this region is at the forefront of LB-mediated cognitive changes.

The control cases were on average younger (65.4 +/- 8.9 years) than those with disease (PD=78.1 +/- 8.7; PDD=76.9 +/- 7.7; DLB=73.2 +/- 6.7 years) (**Table S1**). All four groups were predominantly male and featured similar post-mortem intervals (PMI). The limited number of MMSE scores available proximate to death revealed significant impairment among those with PDD and DLB. Pathological traits available for these brain tissues included levels of global Aβ neuritic plaque and tau NFT deposition as measured by the Consortium to Establish a Registry for Alzheimer’s Disease (CERAD) criteria and the Braak staging system, respectively [27, 28]. Severity levels of LB deposition in the frontal cortex were measured on a semi-quantitative severity scale of 0 to 3. Nearly all demented cases harbored LBs in the frontal cortex, ranging from mild to severe. Amyloid co-pathology was also common among the LBD cases, with half featuring moderate to severe neuritic plaque deposition (CERAD=2-3). In contrast, tau accumulation was sparser. Most LBD cases harbored only mild to moderate NFT levels (Braak I-IV). A small number of PDD and DLB cases (*n*=9) had severe NFT levels (Braak V-IV), increasing the likelihood of clinical symptoms attributable to both LBD and AD-associated neuropathological change [26].

Tandem mass tag mass spectrometry (TMT-MS) quantified 9,661 proteins across all four groups (**Fig. 1A**), including only those proteins quantified in at least 50% of samples. Technical variance was minimized using a tunable median polish approach (TAMPOR), as previously described [29]. The data was then regressed for variability due to age, sex, and PMI. Prior to building a co-expression network, we examined the differential expression in each disease compared to controls. All three Lewy body disorders demonstrated a robust number of significantly altered proteins (*p*<0.05) (**Fig. 1B, Table S2**). PDD featured the greatest amount of differential expression with >3000 proteins significantly altered, including 1,542 increased and 1,530 decreased in disease. DLB harbored nearly 2,000 and PD approximately 1,300 significantly altered proteins compared to controls. Amyloid precursor protein (APP), which has historically correlated strongly to Aβ accumulation in our proteomic datasets [20], was significantly elevated in both PDD and DLB but largely unaltered in PD, consistent with the moderate to severe CERAD scores among the demented cases. SNCA was notably increased in all three Lewy body disorders, reaching significance in PD and PDD and approaching significance in DLB (*p*=0.089). In contrast, microtubule associated protein tau (MAPT) levels were not significantly altered in any of the disease groups compared to controls. APP and SNCA levels correlated significantly with neuropathology measures of neuritic plaque and synuclein deposition, respectively (**Fig. 1C**).

**Figure 1.**
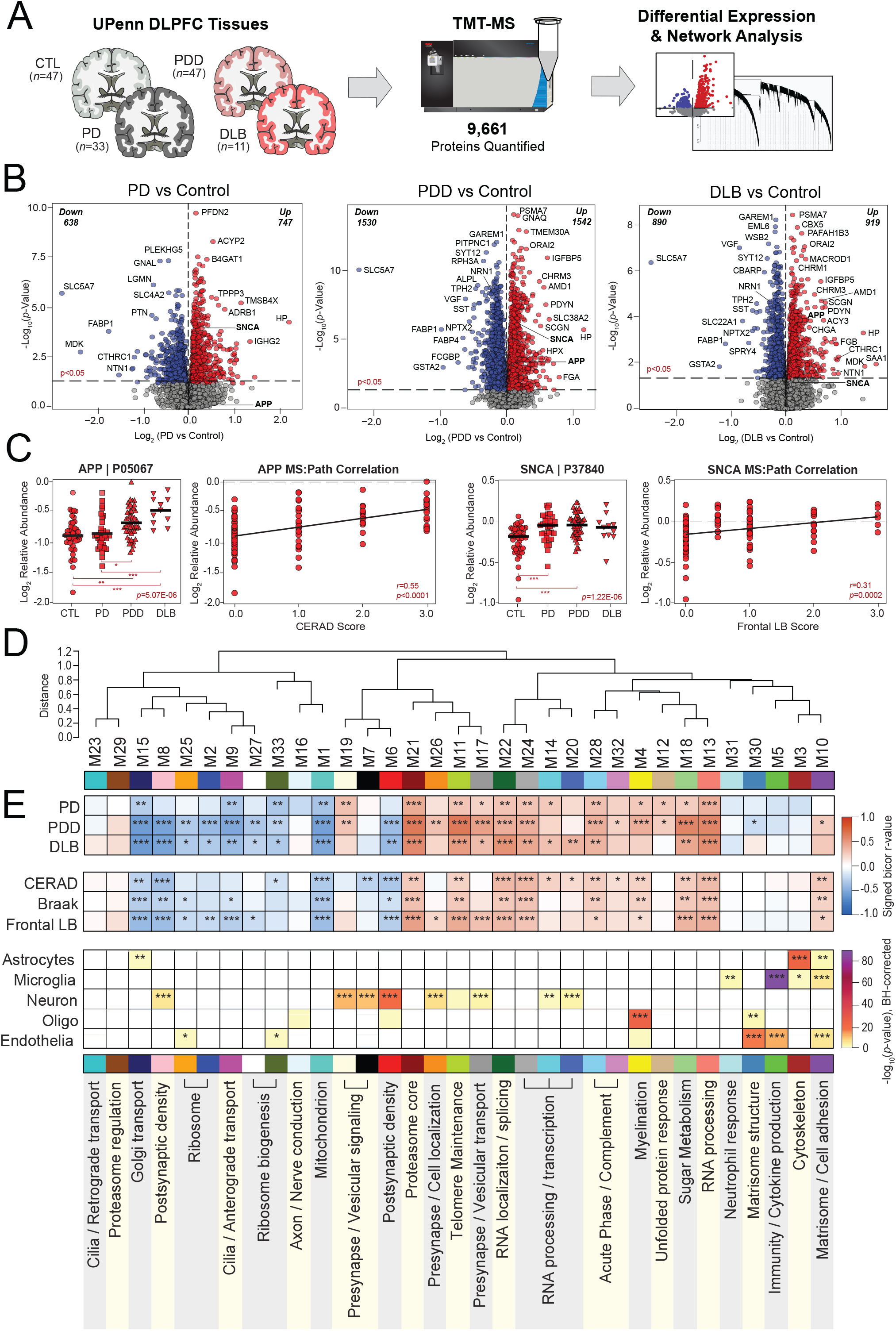
Differential expression and network analysis of UPenn LBD tissues. **(A)** Study approach for analyzing differential expression and co-expression across the UPenn DLPFC tissues, including control (*n*=47), PD (*n*=33), PDD (*n*=47), and DLB (*n*=11) cases. TMT-MS resulted in the quantification of 9,661 proteins across all cases. **(B)** Volcano plots displaying the log_2_ fold change (x-axis) against the -log_10_ statistical *p* value (y-axis) for proteins differentially expressed between pairwise comparisons of each disease to controls. All *p* values across pairwise comparisons were derived by ANOVA with Tukey post-hoc correction. **(C)** Boxplots of MS-measured APP and SNCA levels and their correlations to neuropathology measures of global amyloid plaque (CERAD) and frontal LB deposition, respectively. ANOVA *p* values are provided for each MS abundance plot (*, *p*<0.05; **, *p*<0.01; ***, *p*<0.001), while the Pearson correlation coefficient with associated *p* value is provided for each correlation analysis. **(D)** Co-expression network generated by WGCNA across all UPenn cases, consisting of 33 modules each labeled with a number and color. Module relatedness is shown in the dendrogram. **(E)** Neuropathological trait correlations, cell type marker enrichment, and principal gene ontology for each module. Module abundances were correlated to each disease diagnosis and measures of pathological burden with positive correlations indicated in red and negative correlations in blue. The cell type nature of each module was assessed by module protein overlap with cell type-specific marker lists of astrocytes, microglia, neurons, oligodendrocytes, and endothelia. Gene ontology analysis was used to identify the primary biology reflected by each module. Asterisks in each heat map indicate the degree of statistical significance of the trait correlation or cell type marker enrichment (*, *p*<0.05; **, *p*<0.01; ***, *p*<0.001). Abbreviations: CTL, control; PD, Parkinson’s disease; PDD, Parkinson’s disease dementia; DLB, Dementia with Lewy bodies; DLPFC, dorsolateral prefrontal cortex; TMT-MS, Tandem mass tag mass spectrometry; APP, amyloid precursor protein; SNCA, α-synuclein.

Proteins most highly increased in demented cases included the proteasome subunit PSMA7, calcium channel modulator ORAI2, and various muscarinic cholinergic receptors (CHRM1, CHRM3). Meanwhile, both dementia groups featured starkly decreased levels of known neuroprotective synapse-associated markers VGF and NPTX2, which are consistently decreased in the brains of those with neurodegeneration [30-38]. Neuritin 1 (NRN1), a synaptic protein linked to cognitive resilience in AD, was also significantly decreased in both PDD and DLB [39, 40]. Cholinergic disruption was evidenced by exceptional decreases in the ion channel transporter SLC5A7 across all three diseases, as large as 5-fold lower in DLB compared to controls. This protein, also known as CHT1, mediates presynaptic high-affinity choline uptake in cholinergic neurons for acetylcholine (ACh) synthesis [41], and its dysfunction has been linked to AD in animal models [42, 43]. Yet, several of the most decreased LBD markers had less well-described links to neurodegeneration, such as the kinase regulator GAREM1 and the serotonin synthesizing enzyme TPH2. Overall, these results highlighted significant differential expression across numerous proteins in all three LB diseases, including changes in well-described markers of neurodegeneration.

### Network analysis of the LBD brain reveals alterations across diverse cell types and molecular functions

To examine global systems-based alterations in LBD, we applied Weighted Gene Co-Expression Network Analysis (WGCNA) as previously described [17, 18, 20], which organizes complex proteomic datasets into groups or modules (M) of proteins with similar expression patterns across individual cases [20]. Our resultant LBD co-expression network comprised 8,517 of the total 9,661 proteins. Approximately 12% of the dataset (*n*=1,144 proteins) did not map strongly to a particular module, consistent with the proportions of unassigned proteins we have encountered in previous network analyses [15-20]. These 8,517 proteins were clustered into 33 co-expression modules, with M1 representing the largest module (*n* proteins=601) and M33 the smallest module (*n* proteins=39) (**Fig. 1D, Table S3**). The weighted expression profile, or eigenprotein, of each module was correlated to disease diagnosis and various clinicopathological traits (**Fig. 1E**). In addition, each module was further characterized by cell type enrichment and gene ontology (GO) analyses (**Fig. 1E, Table S4**), as previously described [16-18].

Over two-thirds of the 33 modules correlated significantly to either PDD or DLB or both. These LBD-associated modules reflected a variety of cell type associations, biological ontologies, and cellular compartments. Modules with strongly negative correlations to disease included those linked to mitochondria (M1), Golgi transport (M15), ribosome biogenesis / function (M2, M25, M27, M33), and the postsynaptic density (M6, M8). In contrast, LBD-associated modules with highly positive disease correlations reflected myelination (M4), matrisome / cell adhesion (M10), telomere maintenance (M11), RNA binding / splicing (M13), presynaptic vesicular transport / signaling (M17, M19, M26), sugar metabolism (M18), and proteasome function (M21) (**Fig. 1E**). Module abundance plots across diagnostic groups supported these correlation analyses, with LBD displaying significantly increased levels of positively correlated modules and significantly decreased levels of negatively correlated modules (**Fig. 2A-H**). Most LBD-associated modules were also highly correlated to one or more of the core neuropathologies (**Fig. 1E**). A few modules stood out for their selectively strong correlations to LB burden. For example, two synaptic modules (M17, M26) featured strong positive correlations to LB deposition over other neuropathological traits (**Fig. 1E**).

**Figure 2.**
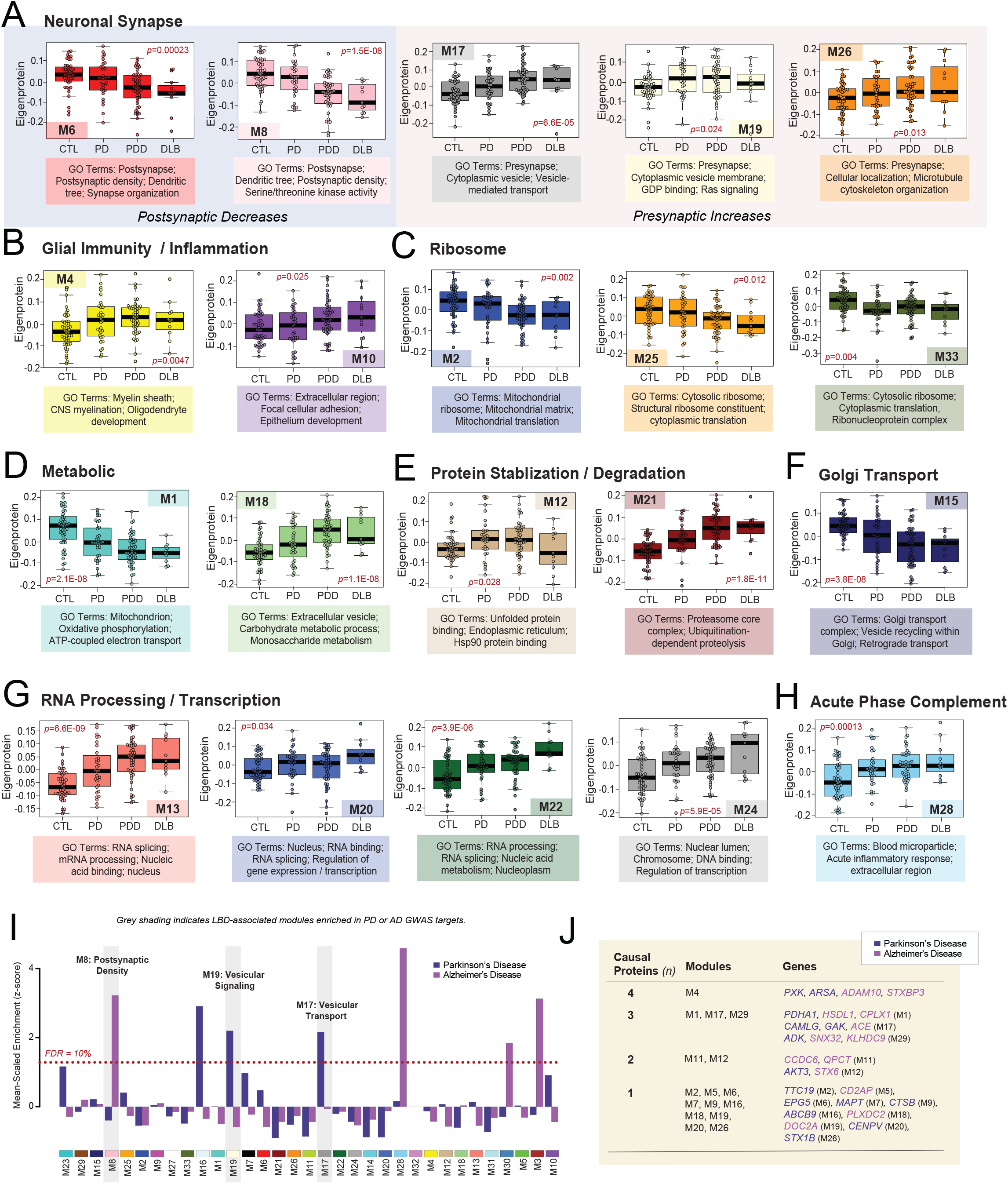
Module abundances of UPenn LBD network reflect significant disease-associated alterations across diverse biological ontologies. **(A-I)** Abundance levels (eigenproteins) of select modules across control and disease groups with their top associated biological ontologies. ANOVA *p* values are provided for each abundance plot. All modules depicted were significantly altered (*p*<0.05) across the four groups. Box plots represent the median and 25th and 75th percentiles, while data points up to 1.5 times the interquartile range from the box hinge define the extent of error bar whiskers. **(J)** Module enrichment of PD (dark blue) and AD (purple) genetic risk factor proteins identified by GWAS. The dashed red line indicates a z score of 1.28, above which enrichment was significant (*p*=0.05) with an FDR of <10%. Gray shading indicates those modules with significant LBD-associated alterations in disease that were also significantly enriched with GWAS targets. Modules are ordered by relatedness as showed in Fig. 1D. **(K)** Table highlighting modules containing proteins identified by integrative multi-omic analysis as maintaining a causal role in PD (dark blue) and AD (purple). Abbreviations: FDR, false discovery rate.

These LB-associated synaptic modules (M17, M26), as well as a third synaptic module (M19), displayed particularly notable abundance trends. In our prior AD networks, synaptic modules were uniformly decreased in disease [17, 18, 20]. Yet, these three synaptic modules demonstrated significant increases in both PDD and DLB (**Fig. 2A**). The presynaptic compartment and related functions (vesicular signaling / transport, cell localization) were most strongly reflected among the top GO terms for these elevated modules (**Table S4**). Accordingly, M19 contained SNCA among its module members, a protein that has been repeatedly linked to presynaptic signaling and membrane trafficking [44]. There was a fourth presynaptic module (M7) that was not significantly altered in LBD, suggesting some selectivity to the upregulation of presynaptic proteins. Meanwhile, neuronal modules more strongly associated with the postsynaptic compartment (M6, M8) were significantly decreased in PDD and DLB, aligning more with our previous observations in AD (**Fig. 2A**). These postsynaptic modules featured strong negative correlations to neuritic plaque, tau, and LB pathology levels (**Fig. 1E**). In sum, these results suggested that network-level changes among synuclein-associated presynaptic proteins may diverge between LBD and AD.

### LBD-associated modules are enriched in genetic risk targets

To investigate causal relationships to disease among our LBD-associated modules, we analyzed each for enrichment of specific disease associated GWAS targets. This analysis was performed using a gene and gene-set analysis tool called MAGMA [45], as previously described [20]. Given LBD shares a significant number of genetic risk factors with both PD and AD [46, 47], we utilized GWAS targets for these two disorders. Two LBD-associated modules (M17, M19) were significantly enriched in PD GWAS targets (**Fig. 2I, Table S5**). Both were presynaptic modules with significant elevations in disease and positive correlations to LB pathology. M16 was also enriched in PD targets, but this axonal module was not significantly altered in our LBD network. Meanwhile, M8 was the only LBD-associated module enriched with AD GWAS targets (**Fig. 2I, Table S6**). This postsynaptic module with significant decreases in LBD stood in contrast to the presynaptic modules enriched in PD GWAS targets, which demonstrated significant increases in LBD. To complement this GWAS enrichment analysis, we also examined our modules for the inclusion of proteins identified in a recent integrative multi-omic analysis as maintaining a pleiotropic or causal role in neuropsychiatric disease, including PD and AD [48]. Of these 48 causal proteins, 27 mapped to a module in our LBD network (**Fig. 2J**). Again, synaptic modules were highly represented among those featuring causal proteins, most notably M17 which harbored PD causal proteins calcium modulating ligand (CAMLG) and cyclin G associated kinase (GAK) and the AD causal protein angiotensin 1 converting enzyme (ACE). The remaining presynaptic modules (M7, M19, M26) each featured one causal protein each. Meanwhile, among postsynaptic modules, M6 harbored the PD causal protein ectopic P-granules 5 autophagy tethering factor (EPG5). Other LBD-associated modules linked to causal proteins included M1 mitochondrion, M4 myelination, M11 telomere maintenance, and M12 unfolded protein response. In summary, these findings highlighted modules with potential causative relationships to disease with select synaptic modules once again emerging as interesting given their enrichment with genetic risk targets.

### Alpha-synuclein protein serves as a bottleneck to LBD-associated presynaptic modules

In co-expression protein networks, there are two predominant categories of centrally important molecules: 1) hub proteins and 2) bottleneck proteins. A hub protein is critical to the structure of its assigned module, harboring large numbers of interactors within its community of co-expressed proteins. Thus, the deletion or removal of a hub from a network is often lethal to cells [49, 50]. In contrast, bottleneck proteins mediate the flow of information between modules, representing key connectors across different communities of co-expressed proteins. The disruption of a protein so critical to module communication could partition the network and similarly cause significant harm to the cell [49, 50]. Given its central neuropathological role in LBD, we were interested in whether SNCA played one or more of these key roles in our proteomic co-expression network.

WGCNA assigns each protein to only one module based on the strength of its correlation to the module eigenprotein [51]. This correlation metric (kME) can also be used to identify module hubs, which are often defined as those proteins ranking among the top 20% of module members based on kME [49, 51]. Using these criteria, SNCA was not a hub of its assigned module (M19). While it featured a moderately strong correlation to M19 (kME=0.6838), SNCA ranked 95 among its 242 module members and fell well outside of hub status (**Fig. 3A**). Yet, we noticed that SNCA harbored kMEs of similar strength to other modules beyond M19, including M7 (kME=0.6245), M17 (kME=0.5820), and M26 (kME=0.5688) (**Table S3**). Like M19, all three of these modules featured strong links to presynaptic gene ontologies. Furthermore, M17 and M26 were also highly correlated to LB deposition, as opposed to neuritic plaque and NFT accumulation. These observations indicated that while it was not a hub of its assigned module, SNCA may play more of a central role in the communication between this group of closely related presynaptic modules.

**Figure 3.**
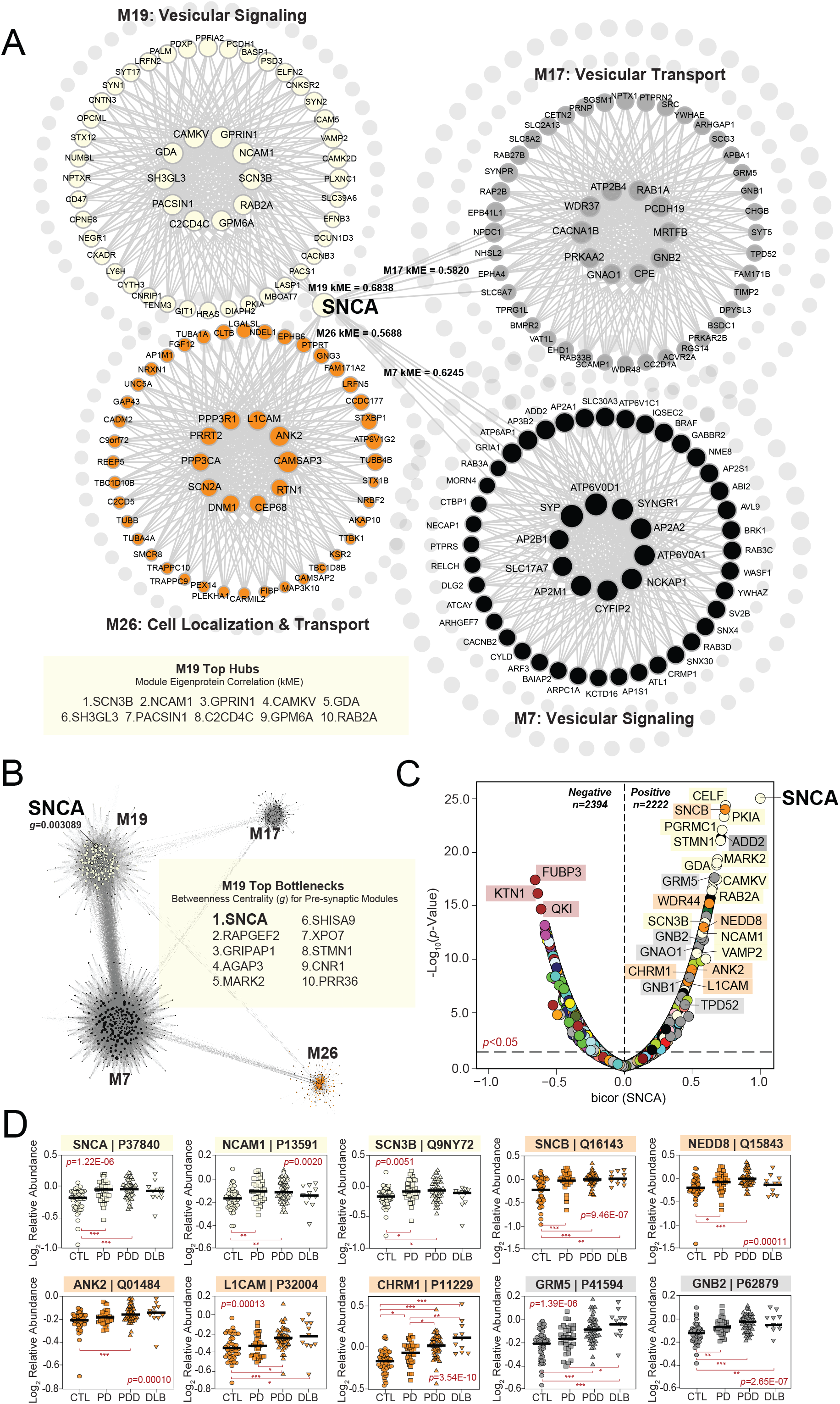
Alpha-synuclein serves as a bottleneck protein between presynaptic UPenn modules. **(A)** Graphical representation of SNCA module membership (kME) relative to four neuronal modules in the UPenn co-expression network. While assigned to M19, SNCA was not a hub of this module. In addition, SNCA maintained moderately strong correlations to M7, M17, and M26 with kME values approaching or exceeding 0.6. All four modules were linked to presynaptic biological ontologies. The top hub proteins for M19 are shown. **(B)** Graphical representation of the bottleneck analysis performed among the four presynaptic modules of interest. Top bottlenecks for M19 across these modules are shown based on measures of betweenness centrality (*g*). SNCA featured the highest g value of M19, indicating its central role in information flow between the four depicted modules. **(C)** Volcano plot displaying the biweight midcorrelation (bicor) to SNCA abundance (x-axis) against the -log_10_ statistical *p* value (y-axis) for all proteins quantified in the UPenn dataset. Proteins are shaded according to color of module membership. There were 2394 proteins with statistically significant (*p*<0.05) negative correlations and 2222 proteins with significant positive correlations to SNCA abundance. Presynaptic modules were among those most highly represented among proteins with the strongest positive SNCA correlations. SNCA -log_10_ *p* value set was from >200 to 25 to keep plot scale. **(D)** Abundance plots for select individual proteins with significant positive correlations to SNCA abundance, highlighting presynaptic modules M17, M19, and M26. ANOVA *p* values are provided for each abundance plot (*, *p*<0.05; **, *p*<0.01; ***, *p*<0.001). Module eigenprotein box plots represent the median and 25th and 75th percentiles, while data points up to 1.5 times the interquartile range from each box hinge define the extent of error bar whiskers. Abbreviations: SNCA, α-synuclein.

We thus investigated whether SNCA might serve as a bottleneck protein between these four modules. In network theory, bottleneck status is typically measured using the metric “betweenness centrality (*g*)”, which in WGCNA can be calculated utilizing the topological overlap matrix generated with each network [51, 52]. Applying this method, we were thus able to measure and rank the top bottleneck proteins among the presynaptic modules of interest. This analysis revealed SNCA as the strongest bottleneck (*g*=0.003089) among the 242 members of M19, highlighting this protein as central to flow of information between the four modules (**Fig. 3B, Table S7**). This result underscored the importance of SNCA within the UPenn LBD network and further bolstered the links we previously observed between these presynaptic modules and neuropathological LB burden.

Accordingly, the same presynaptic modules were also highly represented among individual proteins with strong positive correlations to SNCA abundance levels (**Fig. 3C-D, Table S8**). These highly correlated proteins included known SNCA interactors, such as beta-synuclein (SNCB) of M26 and synaptobrevin-2 (VAMP2) of M19 [44, 53, 54]. M19 and M26 also harbored numerous other SNCA-correlated cell surface proteins, such as L1 cell adhesion molecule (L1CAM), neural cell adhesion molecule 1 (NCAM1), and ankyrin 2 (ANK2). Synuclein levels in L1CAM-positive plasma-derived exosomes have emerged recently as a possible prodromal PD biomarker, supporting the strong association between these proteins [55, 56]. In addition, the highly positive correlation between SNCA and cholinergic receptor muscarinic 1 (CHRM1) of M26 suggested synuclein-mediated cholinergic dysfunction may play a key role in LBD pathophysiology. Many signaling molecules from M17 were also among those proteins highly correlated to SNCA, including various members of the G protein family (GNAO1, GNB1, GNB2, GRM5). Of note, select SNCA-correlated proteins among these presynaptic modules appeared more strongly linked to protein transport and targeting, such as tumor protein D52 (TPD52) of M17 and NEDD8 ubiquitin like modifier (NEDD8) of M26. TPD52 localizes primarily to endoplasmic reticulum (ER), highlighting the complex interactions between the ER and synaptic compartments. Indeed, the ER of neurons is known to mediate several aspects of synaptic transmission, including calcium signaling / homeostasis and vesicular trafficking [57-59].

In sum, we established that SNCA serves as a key bottleneck node to modules with strong links to presynaptic ontologies, robust elevations in LBD, and positive correlations to LB pathology. Numerous individual proteins across these modules demonstrated strong correlations to SNCA abundance and together reflected a wide range of synapse-associated processes.

### LBD-associated network alterations are preserved in replication analyses

#### Emory Replication Analysis

To examine the validity of our UPenn LBD network findings, we analyzed the proteome of a separate cohort of DLPFC tissues derived from the Emory University ADRC brain bank. These cases included tissues with neuropathologically confirmed diagnoses of control (*n*=15), PDD (*n*=10), and DLB (*n*=19). Like the UPenn cohort, those Emory cases with dementia were on average in their mid-70s (PDD=75.3 +/- 10.7, DLB=74.6 +/- 7.9) and predominantly male (**Table S9**). Neuritic plaque deposition was common among the Emory LBD cases. Nearly all Emory DLB cases (*n*=17) and half of the PDD cases (*n*=5) featured moderate to severe levels of plaque deposition (CERAD 2-3). NFT tau levels were overall milder. Yet, 10 of the 19 DLB cases featured severe NFT deposition (Braak V-VI), increasing the likelihood that both AD and LBD pathology were contributing to cognitive decline within this group. All Emory LBD cases featured some degree of LB deposition throughout the frontal cortex. Almost all DLB cases harbored frequent frontal LB inclusions, while those with PDD generally maintained lower burdens often ranging from sparse to moderate.

TMT-MS analysis across all 44 Emory cases quantified 8,213 proteins (**Fig. 4A**), including only those proteins quantified in at least 50% of samples. Like the UPenn cohort, TAMPOR was used to minimize technical variance and the protein abundance data was regressed for variance due to age, sex, and PMI [17, 18, 29]. We then used WGCNA to build a co-expression network on the dataset. The resultant network comprised 7,829 proteins organized into 39 Emory (E) modules with the largest (E-M1) featuring 431 proteins and the smallest (E-M39) harboring 90 proteins (**Fig. 4B, Table S10-11**). Module eigenprotein correlations were similar to those observed in the UPenn LBD network. This included strongly negative disease and pathological correlations among modules linked to postsynaptic (E-M3, E-M10, E-M25), ribosomal (E-M23, E-M36), mitochondrial (E-M2, E-M5, E-M19), and cilia function (E-M6, E-M33). Highly positive disease and pathological correlations among modules linked to the matrisome / cell adhesion (E-M8), proteasome (E-M11), transcription / RNA localization (E-M12), and ER / synaptic signaling (E-M24) also mirrored our UPenn findings. Module preservation analysis revealed that all 39 Emory modules were highly preserved (z score > 10) in the UPenn network (**Fig. 4C**). To directly compare the expression trends of preserved modules across the two LBD networks, we examined the weighted module abundance of the top 20% of proteins by kME in each Emory module within the UPenn dataset. These UPenn “synthetic eigenproteins” revealed strong concordance between abundance trends in both networks across a variety of biological ontologies (**Fig. 4D**).

**Figure 4.**
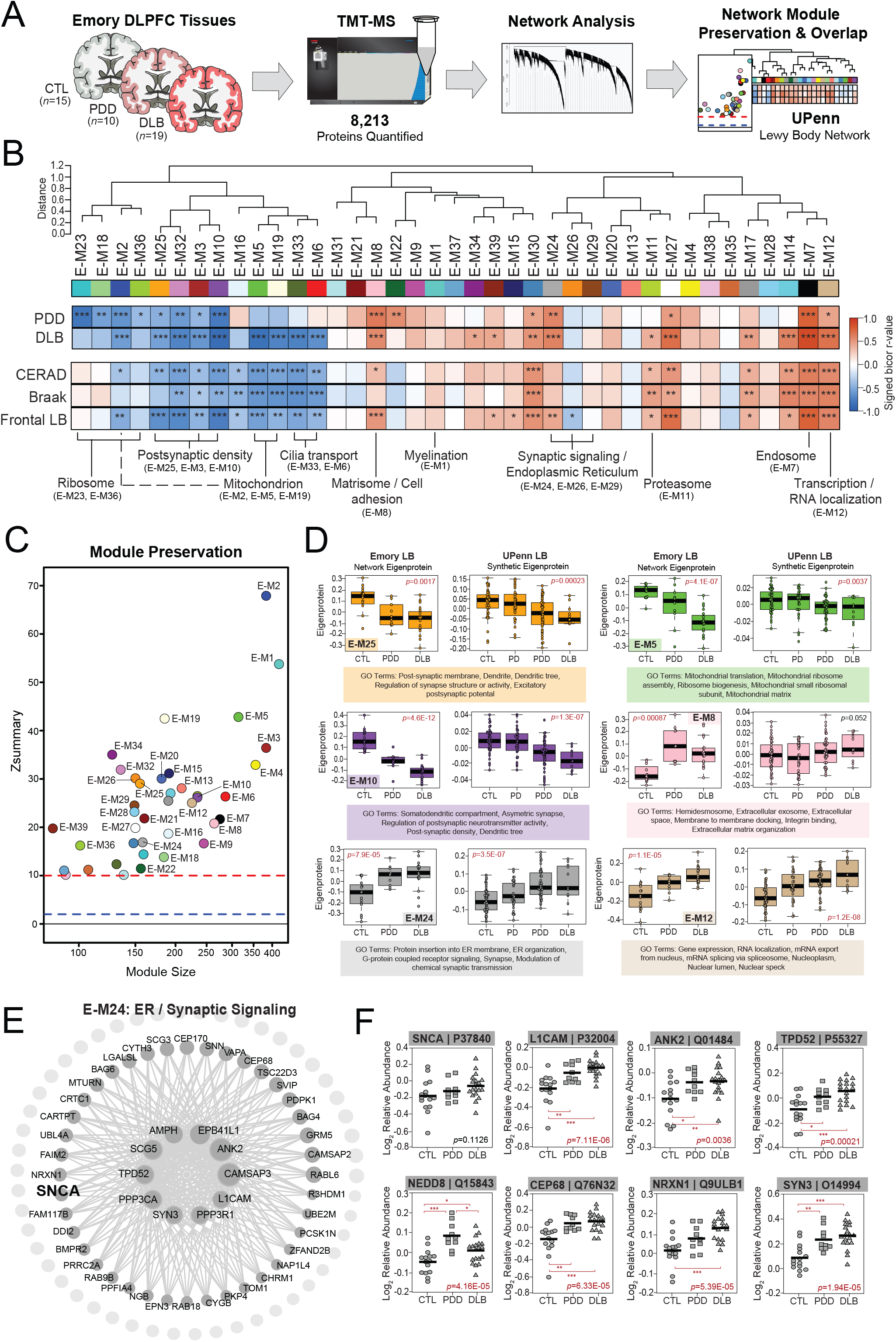
LBD-associated network alterations are replicated in an Emory tissue cohort. **(A)** Study approach for analyzing co-expression across the Emory DLPFC tissues and comparing these network-level alterations to the UPenn dataset. TMT-MS resulted in the quantification of 8,213 proteins across all cases, which included 15 controls, 10 PDD, and 19 DLB tissues. The Emory and UPenn networks were compared using module preservation and overlap analyses. Co-expression network generated by WGCNA across all Emory cases, consisting of 39 modules each labeled with a number and color. Module relatedness is shown in the dendrogram. As in the UPenn network, module abundances were correlated to each disease diagnosis and measures of pathological burden with positive correlations indicated in red and negative correlations in blue. Gene ontology analysis was used to identify the primary biology reflected by each module. Asterisks in each heat map indicate the statistical significance of the trait correlation (*, *p*<0.05; **, *p*<0.01; ***, *p*<0.001). **(C)** Module preservation analysis of Emory network into the UPenn network. Modules with a Z_summary_ score of greater than or equal to 1.96 (*q*=0.05, blue dotted line) were considered preserved, while modules with Z_summary_ scores of greater than or equal to 10 (*q*=1.0E-23, red dotted line) were considered highly preserved. **(D)** Select Emory network module eigenproteins with their corresponding synthetic eigenproteins in the UPenn network. The UPenn synthetic eigenproteins reflected the weighted module abundance of the top 20% of proteins by kME comprising each Emory module. All Emory module eigenproteins shown were significantly altered (*p*<0.05) across groups with synthetic eigenproteins that replicated in the UPenn network. ANOVA *p* values are provided for each eigenprotein plot. Box plots represent the median and 25th and 75th percentiles, while data points up to 1.5 times the interquartile range from the box hinge define the extent of error bar whiskers. **(E)** Graphical representation of individual proteins in the E-24 module arranged by kME with strong hubs at the center. SNCA is designated in bold. **(F)** Plots of individual protein abundances across groups for members of the E24 protein module. ANOVA *p* values are provided for each abundance plot (*, *p*<0.05; **, *p*<0.01; ***, *p*<0.001). Abbreviations: CTL, control; PDD, Parkinson’s disease dementia; DLB, Dementia with Lewy bodies; SNCA, α-synuclein; ER, endoplasmic reticulum.

A hypergeometric Fisher’s exact test (FET) revealed a closely related cluster of Emory modules (E-M24, E-M26, E-M29) that overlapped strongly with the SNCA-associated presynaptic modules in the UPenn network (**Fig. S1**). Accordingly, synaptic signaling ontologies were featured among the top GO terms for these Emory modules (**Table S11**). E-M24 was also strongly linked to ER function, again highlighting the close associations between the synaptic and ER compartments. E-M24 was particularly interesting among this module cluster, as it featured significant increases in LBD and highly positive correlations to frontal LB deposition compared to amyloid and tau. Furthermore, E-M24 harbored SNCA among its module members (**Fig. 4E, Table S10**). As in the UPenn network, SNCA was not a strong hub of E-M24 (kME=0.71). It’s kME ranking of 32 within this relatively small module of 161 proteins placed it just at the edge of the top 20^th^ percentile. SNCA also maintained similar kME values relative to other synapse-associated Emory modules, including E-M26 (kME=0.6481) and E-M29 (kME=0.6210). This suggested a preserved bottleneck role for SNCA within the Emory network as well. In addition to SNCA, E-M24 also featured stark elevations in many of the same synapse-associated proteins also increased in the UPenn network, including L1CAM, ANK2, TPD52, and NEDD8 (**Fig. 4F**). Overall, these results from our Emory cases validated many of the observations from the UPenn network, including increases in proteins associated with SNCA and presynaptic functions.

#### ROSMAP Replication Analysis

As a second replication study, we also examined the proteomes of DLPFC tissues derived from the Religious Orders Study or Rush Memory and Aging Project (ROSMAP) cohorts [60-62]. We used available clinical and pathological traits to classify these cases into control (*n*=42), asymptomatic LB pathology (AsymLB, *n*=21), and Lewy body dementia (LBD, *n*=40) groups (**Table S12**). Control cases comprised those with no cognitive impairment or corticolimbic LB deposition at death. AsymLB was defined as cases with corticolimbic LB deposition but normal cognition at death. Finally, LBD cases included those with clinical dementia and corticolimbic LB pathology present at death. Control cases featured minimal to no AD pathology, while AsymLB and LBD cases were restricted to those with only mild to moderate NFT deposition. These criteria helped ensure a high likelihood that LB deposition, as opposed to AD pathology, was the primary contributor to cognitive decline in our LBD cases [26].

Using TMT-MS, we quantified a total of 7,801 proteins across these cases and built a co-expression network with WGCNA comprising 28 modules. All quantified data was regressed for age, sex, and PMI. We found that the co-expression observed in the UPenn data remained consistent among these ROSMAP cases. All 33 UPenn modules were significantly preserved in the ROSMAP dataset with the majority surpassing preservation z scores of 10 (**Fig. S2A**). Synthetic eigenproteins of UPenn module members within the ROSMAP dataset demonstrated concordance between the two cohorts across multiple modules, including the presynaptic UPenn modules of interest (M17, M19, M26) (**Fig. S2B**). Interestingly, among these presynaptic modules, increased protein levels were also observed in those with AsymLB, suggesting these alterations occur early in preclinical LB deposition. In sum, these results provided additional validation of the disease-associated alterations observed in our discovery UPenn cohort.

### Network level proteome comparison reveals LBD presynaptic co-expression signatures distinct from AD

Unique pathophysiological signatures of LBD that distinguish it from AD and other related dementias are necessary to advance diagnostic and therapeutic target development. Our LBD network observations suggested key differences in co-expression patterns between LBD and our previous AD networks [15-22] that could inform key LBD biomarker discovery. To further compare network-level changes between LBD and AD, we performed a series of overlap analyses with our UPenn LBD network and two separate AD co-expression networks (**Fig. 5A**). The first AD network was also derived from the UPenn ADRC, comprising 49 DLPFC tissues from pathologically defined AD cases and the same 47 control tissues analyzed in the LBD network (**Table S1**). The second AD network was a previously published consensus analysis comprising > 500 control, asymptomatic AD (AsymAD), and AD DLPFC cases from the Banner Sun Health Research Institute [63] and ROSMAP [60-62]. AsymAD cases were defined as those with an Aβ and NFT burden similar to pathologically defined AD cases but without significant cognitive impairment close to death [17]. Thus, these tissues represented an early preclinical phase of AD [64]. We first analyzed the preservation of the 33 modules in our LBD network in these two AD networks (**Fig. 5B-C**). Nearly all LBD modules were highly preserved (Z_summary_ > 10) in both AD networks, indicating that the framework of protein co-expression was consistent across all three datasets.

**Figure 5.**
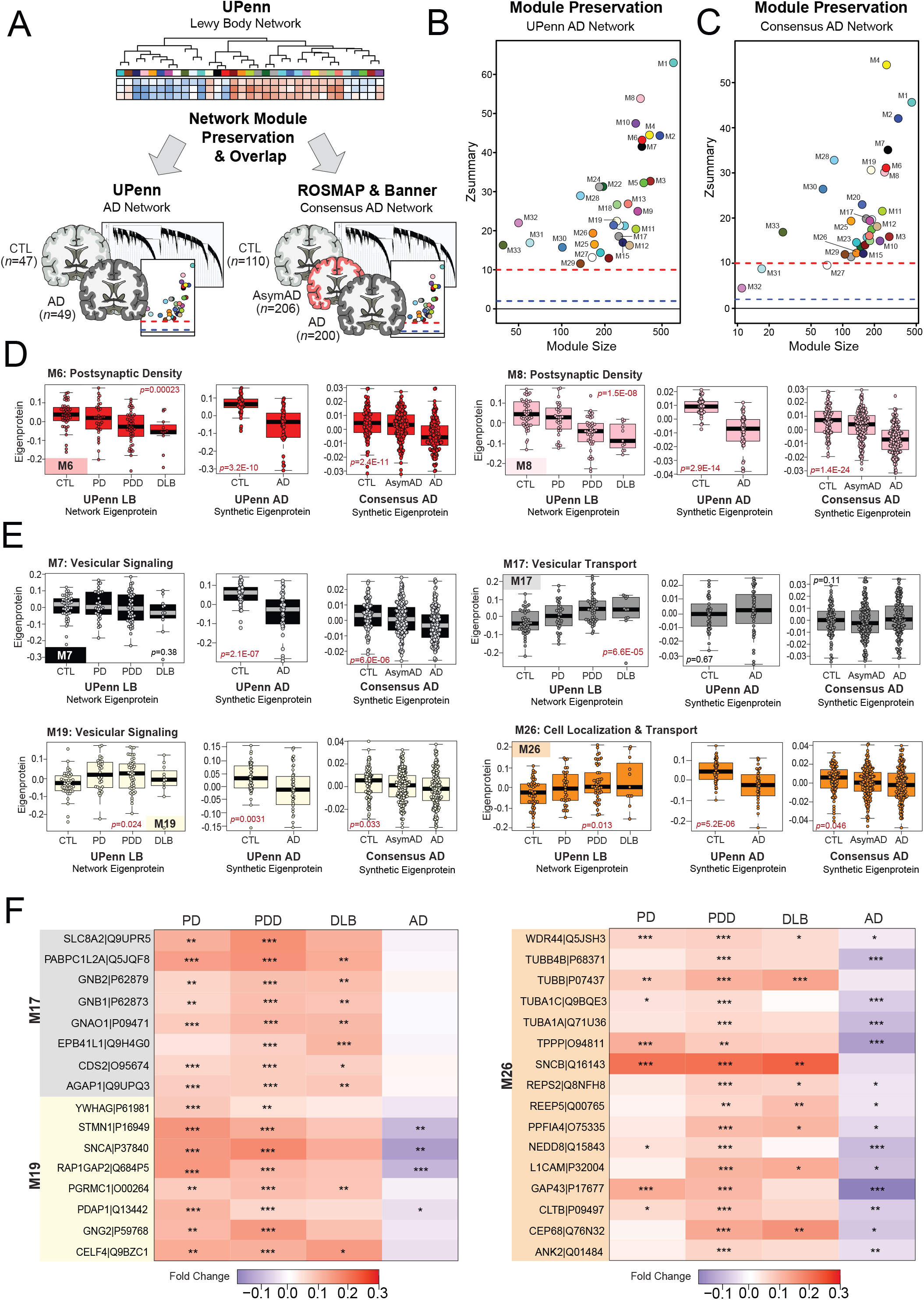
Network level proteome comparison between LBD and AD reveals divergent signatures. **(A)** Study approach for comparing network-level alterations between the UPenn LBD network and two independent AD networks. The first AD network comprised DLPFC UPenn tissues, including 49 AD cases and the same 47 control cases analyzed in the initial UPenn LBD network. The second AD network was a previously published consensus analysis comprising > 500 control, asymptomatic AD (AsymAD), and AD DLPFC cases from the Banner Sun Health Research Institute and Rush Memory and Aging Project (ROSMAP) cohorts. Module preservation and overlap analyses were used to compare these networks. **(B-C)** Module preservation analyses of the UPenn LBD network into the UPenn and Consensus Banner / ROSMAP AD networks. Modules with a Z_summary_ score of greater than or equal to 1.96 (*q*=0.05, blue dotted line) were considered preserved, while modules with Z_summary_ scores of greater than or equal to 10 (*q*=1.0E-23, red dotted line) were considered highly preserved. **(D-E)** UPenn LBD network module eigenproteins associated with the postsynaptic **(D)** and presynaptic **(E)** compartments with their corresponding synthetic eigenproteins in the two AD networks. The AD synthetic eigenproteins reflected the weighted module abundance of the top 20% of proteins by kME comprising each LBD module. ANOVA *p* values are provided for each eigenprotein plot. Box plots represent the median and 25th and 75th percentiles, while data points up to 1.5 times the interquartile range from the box hinge define the extent of error bar whiskers. **(F)** Heat maps depicting the fold change magnitude of select presynaptic proteins across UPenn LBD and AD cases. Increases in protein levels are indicated in red, while decreases are in blue. Asterisks in each heat map indicate the statistical significance of the fold change (*, *p*<0.05; **, *p*<0.01; ***, *p*<0.001). Abbreviations: CTL, control; AsymAD, Asymptomatic Alzheimer’s disease; AD, Alzheimer’s disease; PDD, Parkinson’s disease dementia; DLB, Dementia with Lewy bodies.

To compare the direction of expression of preserved modules across networks, we examined the synthetic eigenproteins of the Lewy body modules in each AD network. As with our above replication analyses, these AD synthetic eigenproteins reflected the weighted expression profiles of the top 20% of proteins in each UPenn LBD module. As expected, synaptic modules significantly decreased in LBD were also significantly decreased in AD (**Fig. 5D**). This included M6 and M8, which were both associated with the postsynaptic density. In contrast, those modules associated with presynaptic functions (M7, M17, M19, M26) showed much less concordance between LBD and AD (**Fig. 5E**). M19 and M26 were significantly increased in LBD but decreased in AD, while M17 featured significant increases in LBD but remained largely unchanged in AD. Finally, M7 was unchanged in LBD but significantly decreased in AD. Individual proteins with the starkest divergence in expression trends between LBD and AD are highlighted in **Fig. 5F**. These differentially expressed markers included SNCA and several co-expressed proteins of interest, such as L1CAM, ANK2, NEDD8, and CEP68. These findings supported our observations that the LBD frontal cortex harbored expression trends among synaptic proteins that diverged from those found in AD. These results also suggested that these two diseases may feature marked differences specifically in presynaptic protein pathophysiology.

### Matrisome-associated protein levels differentiate LBD cases with high levels of amyloid co-pathology

Overlapping AD pathology is extremely frequent in both DLB and PDD with as many as 90% of cases harboring accumulation of the extracellular amyloid-beta (Aβ) plaques [10]. The presence of this concurrent pathology can also influence clinical progression and disease severity. Therefore, we were interested in examining protein expression trends across cases with low and high AD pathology burden. In our UPenn cohort, nearly half of the 58 LBD cases harbored minimal to no Aβ plaque pathology (CERAD 0-1), including 26 PDD and 2 DLB cases. The remaining cases, including 21 PDD and 9 DLB, featured moderate to severe plaque pathology (CERAD 2-3). Differential expression analysis of these low- and high-amyloid LBD cases revealed >700 proteins significantly altered between these two groups (**Fig. 6A, Table S13**). As expected, this included APP and several additional members of M13 (NXPH1, OSTM1), which were significantly elevated in the high-amyloid cases compared to those with low amyloid levels. Yet, most altered among the differentially expressed high- vs low-amyloid markers were numerous members of M10 (MDK, NTN1, SMOC1, CTHRC1), a glial and endothelial enriched module highly linked to the extracellular matrix (i.e., matrisome) and cell adhesion. Compared to low-amyloid LBD cases, these M10 markers demonstrated large, highly significant fold-change elevations in those with high-amyloid LBD and were even more elevated in the UPenn AD cases (**Fig. 6A-B**). These results aligned well with our previous observations in the AD brain network, in which these matrisome proteins serve as hubs of a highly preserved plaque-associated module consistently elevated in AD [17, 18]. Interestingly, certain matrisome markers were significantly decreased in low-amyloid LBD compared to controls (**Fig. 6B-C**), suggesting a separate physiological process in this cohort in which the expression of these plaque-associated proteins is actively suppressed. Such markers included midkine (MDK), a heparin-binding growth factor involved in cell growth and angiogenesis, and collagen triple helix repeat containing 1 (CTHRC1), a protein implicated in vascular remodeling and the cellular response to arterial injury [65, 66]. Both are often elevated in not only AD, but also cancer and tumorigenesis. Overall, these results revealed how the LBD brain network can be a source of not only unique pathophysiological markers, but also those that overlap with other neurodegenerative conditions, distinguishing those LBD cases with mixed pathology from cases with more pure Lewy body deposition. It also highlighted potential divergent glial-mediated pathophysiology in LBD cases with low versus high amyloid plaque burdens.

**Figure 6.**
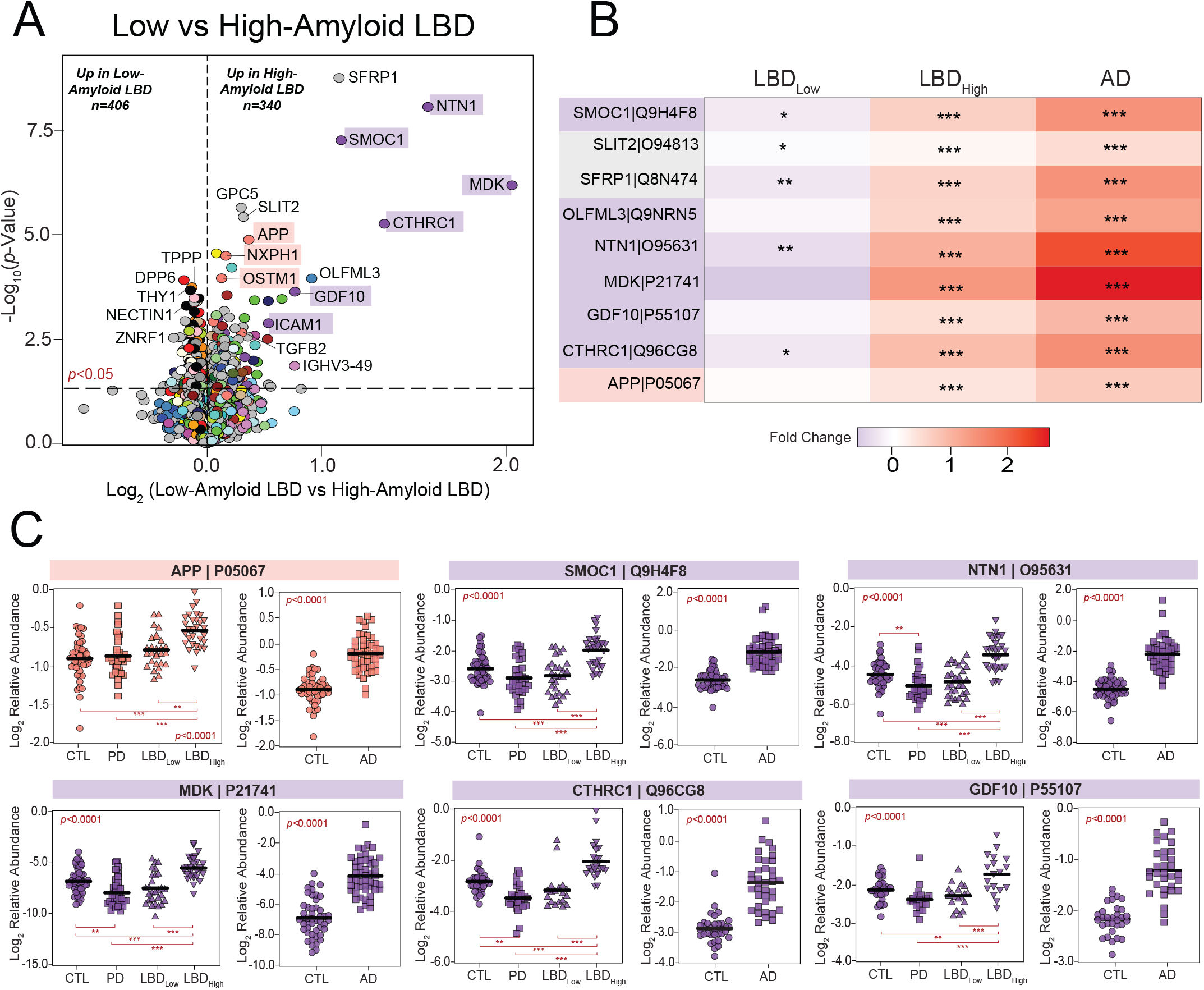
Matrisome proteins distinguish UPenn LBD cases with low and high amyloid burden. **(A)** Volcano plot displaying the log_2_ fold change (x-axis) against the -log_10_ statistical *p* value (y-axis) for proteins differentially expressed between UPenn LBD cases with low (CERAD 0-1) versus high (CERAD 2-3) amyloid deposition. All *p* values were derived by t-test analysis. Proteins are shaded according to color of module membership. Proteins mapping to M10 matrisome in the UPenn LBD network were among those most differentially expressed. **(B)** Heat maps depicting the fold change magnitude of select matrisome and other proteins across UPenn LBD and AD cases. Increases in protein levels are indicated in red, while decreases are in blue. Asterisks in each heat map indicate the statistical significance of the fold change (*, *p*<0.05; **, *p*<0.01; ***, *p*<0.001). (C) Plots of individual protein abundances across groups, including low- and high-amyloid LBD, in the UPenn LBD and AD datasets. ANOVA *p* values are provided for each abundance plot (*, *p*<0.05; **, *p*<0.01; ***, *p*<0.001). Abbreviations: CTL, control; PD, Parkinson’s disease; LBD_Low_, Low-amyloid Lewy body dementia; LBD_High_, High-amyloid Lewy body dementia.

## Discussion

The corticolimbic pathophysiology underlying the aggressive cognitive and neuropsychiatric deterioration in LBD is extremely complex, poorly understood, and features significant overlap with AD. In the current study, we employed co-expression network proteomics to define systems-based pathophysiologic alterations in the frontal cortex of a large UPenn autopsy cohort and compare these signatures to those observed in AD. We identified a diverse array of protein modules altered in the brains of those with PDD and DLB, encompassing synaptic, metabolic, and inflammatory pathophysiology. We then validated these network signatures across independent LBD cohorts and identified reproducible synaptic alterations that diverged from those in the AD brain. We also identified informative overlapping signatures between LBD and AD, including glial-associated matrisome markers that proved highly concordant with Aβ deposition and capable of stratifying LBD cases with low versus high burdens of amyloid plaque co-pathology. These results underscore how proteomic co-expression network analysis can yield insights into key divergent and overlapping pathophysiological signatures in the LBD and AD brain.

Synaptic protein loss is generally considered a universal feature of the neuropathological changes observed in dementia. Numerous studies in AD have shown that pathological measures of synaptic loss correlate more strongly with cognitive impairment compared to Aβ and tau pathology [67-70]. Accordingly, we and others have observed stark decreases in a variety of synaptic proteins in the AD brain across multiple independent cohorts and brain regions [15-21, 71]. We have also identified these synaptic decreases in the brains of those with AsymAD, or individuals with significant neuritic plaque and NFT deposition but no evidence of clinical cognitive impairment at death [17, 18, 20]. These results suggest early synaptic losses in AD independent of clinical declines. Aligning with these observations, we found significant decreases among our LBD cases in two large modules linked to postsynaptic function (M6, M8). These modules included synaptic markers that already feature well-described decreases in neurodegeneration, such as VGF and NPTX2 [30-38], reinforcing their potential as reliable markers of degeneration across different neurologic diseases.

Yet, we also observed increased levels in presynaptic modules among our UPenn LBD cases, contrasting with our prior AD observations. These included M17, a neuron-enriched module linked to synaptic vesicular transport that featured various GTPases among its hub proteins (RAB1A, GNB2, GNAO1), as well as M19, another neuronal module that included SNCA and other proteins linked to vesicular signaling. The third neuronal module significantly increased in LBD was M26, which maintained ontological associations to both the synapse and cellular localization. While its hub proteins included largely cell surface proteins (ANK2, L1CAM), this module also featured proteins involved in protein targeting, folding, and processing (CEP68, NEDD8). All three of these modules were either unchanged or significantly decreased when examined in our comparison AD networks, indicating divergent pathophysiology between the two dementias. It is unclear why LBD features these presynaptic increases and will require further investigation. On one hand, it is possible these increases represent a compensatory, rather than pathological, response in this brain region. However, the network-based links these modules shared with SNCA levels and LB deposition suggests that aberrant SNCA function is to some extent mediating these findings.

Accordingly, SNCA served as a strong bottleneck node among these presynaptic modules. While hub proteins, which demonstrate high connectivity within their respective modules, are often considered key molecular drivers in co-expression networks [72], there is also growing recognition that proteins with a high degree of connectivity between modules are also highly relevant to disease. These bottleneck proteins have been shown to play central roles in various disorders and serve as successful therapeutic targets [49, 73-75]. As a strong presynaptic bottleneck, SNCA mediated communication between the three modules highlighted above (M17, M19, M26), as well as a fourth larger module (M7) also heavily associated with vesicular signaling and the presynaptic compartment. This aligns with the growing amount of research indicating SNCA localizes to the presynaptic terminal and participates in vesicular cycling, including regulation of vesicle pool size, mobilization, and endocytosis [44]. In addition, these presynaptic modules featured proteins known to interact closely with SNCA, such as synaptobrevin-2 (VAMP2) and synapsin 1 (SYN1) [44, 54, 76]. L1 cell adhesion molecular (L1CAM), an increasingly studied neuronal surface marker for isolating and measuring exosome-associated SNCA in biofluids [55, 56], also mapped to these presynaptic modules (M26) and its increasing abundance in LBD correlated strongly with those of SNCA. Thus, our LBD network underscored well-established links between SNCA and the presynaptic compartment and supported its central role in the unique pathophysiological changes of LBD.

These presynaptic LBD signatures also supported unique alterations in cholinergic pathways and their potential therapeutic implications. M26 featured muscarinic cholinergic receptors CHRM1 (M1) and CHRM4 (M4), which have both demonstrated promise as synaptic targets for cognitive and behavioral symptoms in dementia [77, 78]. Drugs that enhance synaptic M1 and M4 activity are currently being explored in the management of both AD and DLB. Compared to controls, both receptors demonstrated significant increases among LBD cases but stable, largely unchanged levels in AD. While this reveals potential differences in AD and LBD cholinergic function, it does suggest both diseases are able to preserve these receptors to some extent and may be responsive to M1 and M4 agonists. Yet, the stark increases LBD demonstrated in these cholinergic receptors, as well as others (CHRM2, CHRM3), could be in response to significant losses in other components of the cholinergic pathways. For instance, all three LB disorders in our UPenn network demonstrated dramatic, several-fold decreases in SLC5A7, a protein necessary for presynaptic choline uptake and ACh synthesis [41]. Of note, though much of its activity is localized to the presynaptic terminal, SLC5A7 expression trends most closely aligned with postsynaptic M8. Thus, despite their predominant ontologies, it is important to note that all our neuronal modules likely harbor some mixture of proteins that function in both the presynaptic and postsynaptic spaces. Nevertheless, these cholinergic trends further showcase the diversity of abundance alterations within LBD synaptic pathways, highlighting those proteins with preserved to increased levels in disease that may respond well to therapeutic agonists.

Yet, synaptic pathways were not the only ones implicated as distinctly altered in LBD. In both the UPenn and Emory LBD networks, proteins involved in protein targeting, folding, and ER function were also co-expressed with our elevated presynaptic LBD markers. M26 best reflected this co-expression of synaptic and protein processing molecules in the UPenn network, while E-M24 of the Emory network mapped strongly to both synaptic and ER ontologies. These network associations likely reflect the well-established functional relationships between ER and synaptic regulation [57-59]. Many have also linked SNCA itself to ER stress and aberrant protein processing. For instance, Colla et al. found oligomeric species of SNCA in the ER of both animal and human brains with synucleinopathy [79], suggesting an ER-mediated stress response may play an integral role in disease pathophysiology. Others have also reported colocalization of ER stress markers with synuclein inclusions in diseased brain tissue [80, 81]. Furthermore, LB disease has been linked to various disruptions in protein processing, such as neddylation [82]. NEDD8, a ubiquitin-like protein involved prominently in neddylation, was among the synapse-associated modules in our UPenn and Emory networks that demonstrated strong increases in LBD and marked decreases in AD. Dysfunction of this protein has been previously linked to multiple neurodegenerative diseases with ubiquitinated inclusions, including AD and LB disorders [82-84]. However, our results suggest the pathophysiological mechanism underlying this dysfunction may differ between these two disorders.

These divergent network changes could eventually yield much needed diagnostic and therapeutic markers of LBD. The observation in our ROSMAP cases that these presynaptic proteins are elevated in asymptomatic disease further supports their potential clinical utility as early targets. Yet, tools that help identify overlapping LBD and AD are also useful. Studies have shown that the burden and distribution of AD pathology can have a significant impact on LBD presentation and progression [85, 86]. Thus, understanding the physiological breadth of co-pathology among these individuals is also paramount. Our network analysis revealed that proteins associated with the extracellular matrix (ECM) were best at distinguishing LBD cases with low versus high levels of amyloid co-pathology. This supports growing evidence that these proteins associate strongly with amyloid plaques [71]. Furthermore, these results underscore our prior AD brain network analyses, which have established many of these matrisome proteins (SMOC1, NTN1, MDK) as hubs of a highly preserved glia-associated module consistently elevated in both AsymAD and AD [16-18]. Our current LBD data provides an additional role for these emerging biomarkers as early indicators of AD co-pathology in LB populations, which could help guide clinical management throughout the course of disease. In addition, their elevation in such a sizeable portion of LBD patients suggests these markers could provide another avenue independent of SCNA for therapeutic targeting.

Prior integrative proteomic studies from our group and others have demonstrated the translation potential of brain network analysis into promising biofluid markers of disease [16, 18, 23, 71, 87]. We have previously demonstrated up to 70% overlap between the brain and CSF proteomes using TMT-MS. This has allowed us to identify panels of CSF biomarkers in AD reflecting a diverse range of pathophysiology and further validate these markers in cross-sectional and longitudinal AD cohorts [16, 18, 23, 24, 87]. In a recent manuscript, we showed that a panel of 48 CSF AD markers, originally identified in brain network analyses, improved early diagnostic and predictive assessments of sporadic AD [23]. A separate study demonstrated that CSF levels of matrisome markers SMOC1 and SPON1 were elevated nearly 30 years prior to the onset of symptoms in an autosomal dominant AD population [24]. Thus, future directions include integrating the LBD brain network proteome with the CSF and plasma proteomes of diseased patients to similarly identify molecularly diverse biofluid panels that could advance the diagnostic and predictive accuracy of LBD.

Our focus on the DLPFC could be viewed as a limitation in this study. This region was chosen because it is commonly affected in diffuse neocortical LBD and routinely scored in its neuropathological diagnosis. In addition, frontal executive deficits are commonly among the first symptoms observed in LBD [26], indicating this region could provide a valuable map of early pathophysiological changes in the evolution of LB-mediated cognitive changes. The robust differential expression we observed even in non-demented PD cases further suggests this region is affected early in the brainstem-to-corticolimbic disease evolution thought to eventually provoke PDD. Yet, it is possible other heavily affected regions in LBD, including the inferior temporal and parietal lobes, may yield different network findings and additional insights. Among other limitations, our LBD subjects lacked racial diversity and skewed predominantly male. In our prior network studies of the AD brain proteome, we have found that both age and sex have a very limited impact on disease-associated module trends [18]. Yet, it will be important in future studies, particularly when examining these markers in biofluids, to utilize large, more diverse cohorts to define the impact of these demographic variables on LBD protein signatures.

In summary, our study offers a network-level map of the LBD brain proteome, which revealed disease-associated alterations in a diverse range of protein systems. Using this network, we were able to identify key overlapping and divergent protein signatures in LBD and AD tissues and correlate these disease-associated alterations to core neuropathologies. These results can serve as a strong systems-based framework for future integrative studies focused on identifying protein biofluid markers relevant to corticolimbic pathophysiology in the LBD brain.

## Methods

### Brain Tissues

Human postmortem brain tissues used in this study were obtained from the UPenn ADRC, Emory ADRC, and ROSMAP [60-62] autopsy collections. All tissues were derived from the DLPFC (BA 9) and acquired under Institutional Review Board protocols at each representative institution. All ROSMAP participants signed informed and repository consents and an Anatomic Gift Act. In all cases, neuritic plaque distribution was scored according to the Consortium to Establish a Registry for Alzheimer’s Disease (CERAD) criteria [28], and the extent of neurofibrillary tangle pathology was assessed with the Braak staging system [27]. The frequency of Lewy body (LB) deposition in the frontal cortex was scored in the UPenn and Emory cases using similar semi-quantitative scales. The UPenn scale included scores of 0 (absent), 1 (sparse), 2 (moderate), and 3 (frequent), while the Emory scale comprised scores of 0 (absent), 1 (sparse to moderate), and 2 (frequent). LB deposition in the ROSMAP cases was scored using a global regional scale that indicated whether these inclusions were absent, nigral-predominant, limbic-type, or neocortical-type, as previously described [6]. Clinical scores were available for the UPenn and ROSMAP samples, including Mini-Mental Status Examination (MMSE) values. Clinical consensus cognitive diagnoses were also provided for ROSMAP cases based on detailed neuropsychological testing, which indicated whether an individual had no cognitive impairment (NCI), mild cognitive impairment (MCI), or dementia at death, as previously reported [88-90]. Disease classification in the UPenn and Emory cohorts reflected neuropathological diagnoses provided by expert pathologists at each institution. These diagnoses were made in accordance with established criteria and guidelines [26, 91]. Disease classification of ROSMAP tissues was determined using available clinical and pathological traits. ROSMAP controls included those with NCI, absent corticolimbic LBs, and minimal neuritic plaque and NFT deposition (CERAD 0-1, Braak NFT 0-II). Asymptomatic LB (AsymLB) cases were those with NCI, present corticolimbic LBs, and mild to moderate NFT deposition (Braak NFT 0-IV), while those with Lewy body dementia (LBD) featured dementia at death, present corticolimbic LBs, and mild to moderate NFT deposition (Braak NFT 0-IV). These criteria surrounding tau levels helped ensure a high likelihood that LB deposition, as opposed to AD pathology, was the primary contributor to cognitive decline in our LBD cases [26]. All sample metadata are provided in https://www.synapse.org/#!Synapse:syn53177242. The subsequent Methods sections outline the processing, TMT labeling, and MS analysis of the three tissue cohorts included in this study. All procedures were performed within the Emory University Center for Neurodegenerative Disease and remained largely consistent across cohorts with minor differences where indicated. The 187 UPenn samples included in this study were processed and MS quantified within a larger UPenn tissue cohort comprising a total of 354 samples. Only samples with neuropathologically confirmed diagnoses of control (*n*=47), PD (*n*=33), PDD (*n*=47), DLB (*n*=11), and AD (*n*=49) were included in the subsequent network analyses. Likewise, the 103 ROSMAP cases included for validation were quantified as part of a larger cohort of ROSMAP cases that comprised a total 610 samples. Only those meeting clinical and pathological criteria for control (*n*=42), AsymLB (*n*=21), and LBD *(n*=40) were used for validation analyses. Detailed methods have been previously published for the processing and MS analysis of the complete ROSMAP cohort [92], which we refer to when appropriate.

### Brain Tissue Homogenization and Protein Digestion

Tissue homogenization of all cases was performed essentially as described [20, 93]. Approximately 100 mg (wet weight) of each tissue sample was homogenized in 500 μL of 8 M urea lysis buffer (8 M urea, 10 mM Tris, 100 mM NaH2PO4, pH 8.5) with HALT protease and phosphatase inhibitor cocktail (ThermoFisher). Tissues were added to the lysis buffer immediately after excision in Rino sample tubes (NextAdvance) supplemented with ∼100 μL of stainless-steel beads (0.9 to 2.0 mm blend, NextAdvance). Using a Bullet Blender (NextAdvance), tissues were then homogenized at 4 °C with 2 full 5 min cycles. The lysates were transferred to new Eppendorf Lobind tubes and sonicated for 3 cycles, each lasting 5 seconds at 30% amplitude. Sample lysates were then centrifuged for 5 min at 15,000 x g and the supernatant transferred to new tubes. Protein concentration was determined by bicinchoninic acid (BCA) assay (Pierce). For protein digestion, 100 μg of each sample was aliquoted and volumes normalized with additional lysis buffer. Samples were reduced with 1 mM dithiothreitol (DTT) at room temperature for 30 min followed by 5 mM iodoacetamide (IAA) alkylation in the dark for another 30 min. Lysyl endopeptidase (Wako) at 1:100 (w/w) was added, and digestion allowed to proceed overnight. Samples were then 7-fold diluted with 50 mM ammonium bicarbonate. Trypsin (Promega) was added at 1:50 (w/w) and digestion was carried out for another 16 hours. The peptide solutions were acidified to a final concentration of 1% (vol/vol) formic acid (FA) and 0.1% (vol/vol) trifluoroacetic acid (TFA) before desalting with a 30 mg HLB column (Oasis). Prior to sample loading, each HLB column was rinsed with 1 mL of methanol, washed with 1 mL 50% (vol/vol) acetonitrile (ACN), and equilibrated with 2×1 mL 0.1% (vol/vol) TFA. Samples were then loaded onto the column and washed with 2×1 mL 0.1% (vol/vol) TFA. Elution was performed with 2 volumes of 0.5 mL 50% (vol/vol) ACN. An equal amount of peptide from each sample was aliquoted and pooled as the global internal standard (GIS), a fraction of which was TMT labeled and included in each batch as described below.

### Isobaric Tandem Mass Tag (TMT) Peptide Labeling

TMT peptide labeling was performed as previously described [20, 93]. As outlined above, the 187 UPenn samples included in this study were labeled and MS analyzed within a larger UPenn tissue cohort comprising a total of 354 samples. Prior to labeling, these 354 UPenn cases were randomized into 24 batches by age, sex, and diagnosis. Labeling was performed using TMTpro 16-plex kits (ThermoFisher 44520). Each batch included one TMT channel with a labeled GIS standard. Labeling of sample peptides was performed as previously described [19, 93, 94]. Briefly, each sample (100 μg of peptides) was re-suspended in 100 mM triethylammonium bicarbonate (TEAB) buffer (100 μL). TMT labeling reagents (5 mg) were equilibrated to room temperature. Anhydrous ACN (256 μL) was added to each reagent channel. Each channel was then gently vortexed for 5 minutes. A volume of 41 μL from each TMT channel was transferred to each peptide solution and allowed to incubate for 1 hour at room temperature. The reaction was quenched with 5% (vol/vol) hydroxylamine (8 μL) (Pierce). All channels were then dried by SpeedVac (LabConco) to approximately 150 μL, diluted with 1 mL of 0.1% (vol/vol) TFA, and acidified to a final concentration of 1% (vol/vol) FA and 0.1% (vol/vol) TFA. Labeled peptides were desalted with a 200 mg C18 Sep-Pak column (Waters). Prior to sample loading, each Sep-Pak column was activated with 3 mL of methanol, washed with 3 mL of 50% (vol/vol) ACN, and equilibrated with 2×3 mL of 0.1% TFA. After sample loading, each column was washed with 2×3 mL 0.1% (vol/vol) TFA followed by 2 mL of 1% (vol/vol) FA. Elution was performed with 2 volumes of 1.5 mL 50% (vol/vol) ACN. The eluates were then dried to completeness by SpeedVac. The 44 Emory samples were randomized by age and diagnosis into 3 batches and labeled using TMTpro 16-plex kits (ThermoFisher 44520). Each batch included one TMT channel with a labeled GIS standard. Labeling of these sample peptides then proceeded according to the protocols above. Randomization and multiplex labeling of ROSMAP cases were performed according to very similar protocols, as previously described in detail [92].

### High-pH Off-line Fractionation

High pH fractionation of all cases was performed essentially as described [93, 95] with slight modifications. Dried samples were resuspended in high pH loading buffer comprising 0.07% (vol/vol) NH4OH, 0.045% (vol/vol) FA, and 2% (vol/vol) ACN. Resuspended samples were then loaded onto a Water’s Ethylene Bridged Hybrid (BEH) column (1.7 um, 2.1 mm x 150 mm). A Thermo Vanquish high-performance liquid chromatography (HPLC) system was used to carry out the fractionation. Solvent A consisted of 0.0175% (vol/vol) NH4OH, 0.01125% (vol/vol) FA, and 2% (vol/vol) CAN. Solvent B comprised 0.0175% (vol/vol) NH4OH, 0.01125% (vol/vol) FA, and 90% (vol/vol) ACN. The sample elution was performed over a 25 min gradient with a flow rate of 0.6 mL/min. A total of 192 individual equal volume fractions were collected across the gradient and subsequently pooled by concatenation into 96 fractions [95]. The fractions were then dried to completeness using a SpeedVac.

### Mass Spectrometry Analysis of UPenn Samples

MS analysis was performed on the fractionated UPenn samples as previously described with modifications [17, 18, 20, 93]. Briefly, fractions were resuspended in an equal volume of loading buffer (0.1% FA, 0.03% TFA, 1% ACN) and analyzed by liquid chromatography coupled to tandem mass spectrometry (LC-MS/MS). Peptide eluents were separated on a custom in-house packed Charged Surface Hybrid (CSH) column (1.7 um, 15 cm × 150 μM internal diameter) by a Dionex RSLCnano ultra-performance liquid chromatography (UPLC) system (ThermoFisher Scientific). Buffer A comprised water with 0.1% (vol/vol) FA, and buffer B comprised 80% (vol/vol) ACN in water with 0.1% (vol/vol) FA. Elution was performed over a 30 min gradient with a flow rate of 1500 nL/min. The gradient ranged from 1% to 99% solvent B. Peptides were monitored on a Orbitrap Eclipse mass spectrometer with high-field asymmetric waveform ion mobility spectrometry (FAIMS) (FAIMS Pro Interface, ThermoFisher Scientific). Two compensation voltages were chosen for FAIMS. For each voltage (-45 and -65) top speed cycle of 1.5 seconds, the full scan (MS1) was performed with an m/z range of 410-1600 and 60,000 resolution at standard settings. The higher energy collision-induced dissociation (HCD) tandem scans were collected at 35% collision energy with an isolation of 0.7 m/z, resolution of 30,000 with TurboTMT, AGC setting of 250% normalized AGC target, and a maximum injection time of 54 ms. For all batches, dynamic exclusion was set to exclude previously sequenced peaks for 20 seconds within a 10-ppm isolation window.

### Mass Spectrometry Analysis of Emory Samples

MS analysis on the fractionated Emory samples was performed similarly to the UPenn samples with modifications. Fractions were resuspended in an equal volume of loading buffer (0.1% FA, 0.03% TFA, 1% ACN) prior to LC-MS/MS analysis. Peptide eluents were separated on a custom in-house packed Charged Surface Hybrid (CSH) column (1.7 um, 15 cm × 150 μM internal diameter) by a Dionex RSLCnano running capillary flow UPLC (ThermoFisher Scientific). Buffer A comprised water with 0.1% (vol/vol) FA, and buffer B comprised 80% (vol/vol) ACN in water with 0.1% (vol/vol) FA. Elution was performed over a 40 min gradient with flow rate of 7 uL/min. The gradient ranged from 1% to 99% solvent B. Peptides were monitored on a Orbitrap Exploris 240 mass spectrometer at 2 seconds top speed cycle. Each cycle comprised of a full scan (MS1) with an m/z range of 410-1600 and 120,000 resolution at standard settings. The HCD tandem scans were collected at 36% collision energy with an isolation of 0.7 m/z, resolution of 45,000, AGC setting of 250% normalized AGC target, and 100 ms maximum injection time. For all batches, dynamic exclusion was set to exclude previously sequenced peaks for 20 seconds within a 10-ppm isolation window.

### Mass Spectrometry of ROSMAP Samples

MS analysis of fractionated ROSMAP samples was performed in two separate sets as previously described [92].

### Database Searches and Protein Quantification

All RAW files acquired from TMT-MS of all cases were searched against a human reference protein database using the Proteome Discoverer suite (version 2.4, ThermoFisher Scientific). MS2 spectra were searched against the UniProtKB human proteome database containing Swiss-Prot human reference protein sequences (20,338 target proteins downloaded in 2019). Searches were performed using previously published protocols [17, 18, 93]. Percolator was used to filter peptide spectral matches (PSMs) and peptides to a false discovery rate (FDR) of less than 1%. Following spectral assignment, peptides were assembled into proteins and were further filtered based on the combined probabilities of their constituent peptides to a final FDR of 1%. A multi-consensus in Proteome Discoverer was then performed to achieve parsimonious protein grouping across both sets of samples. In cases of redundancy, shared peptides were assigned to the protein sequence in adherence with the principles of parsimony. As default, the top matching protein or “master protein” was the protein with the largest number of unique peptides and smallest value in the percent peptide coverage (i.e., the longest protein). Reporter ions were quantified using an integration tolerance of 20 ppm with the most confident centroid setting. Only parsimonious peptides were considered for quantification.

### Controlling for Batch-Specific Variance

A tunable median polish approach (TAMPOR) [29] was used to remove technical batch variance in the proteomic data from all three cohorts, as previously described [17]. TAMPOR is utilized to remove inter-batch variance while preserving meaningful biological variance in protein abundance values, normalizing to the median of selected intra-batch samples and the median samplewise abundance, alternately and iteratively in a median polish [29]. We have previously applied this batch-correction approach to multiple large proteomic datasets [17, 18, 92]. This approach is robust to outliers and up to 50% of measurements missing. If a protein had more than 50% of samples with missing values, it was removed from the protein abundance matrix. No imputation of missing values was performed for any cohort. For the current data, TAMPOR leveraged the median protein abundance from the pooled GIS TMT channels as the denominators in both factors to normalize sample-specific protein abundances across batches.

### Regression of Covariates

Following TAMPOR batch correction, the protein abundance matrices from all three cohorts were subjected to non-parametric bootstrap regression by subtracting the covariate of interest multiplied by the median estimated coefficient from 1000 iterations of fitting for each protein in the log_2_(abundance) matrix, as previously described [17]. The UPenn and Emory datasets were regressed for age, sex, and PMI. The ROSMAP cases were regressed for these three covariates, as well as any residual variation related to batch. Ages at death used for regression were uncensored. Case diagnosis was also explicitly modeled and protected in each iteration.

### Weighted Gene Co-expression Network Analysis (WGCNA)

The WGCNA algorithm was used to perform co-expression network analysis on the batch-corrected and regressed data from all three cohorts, as previously described [17, 18, 20]. A total of four co-expression networks were built for this study. For UPenn cases, two separate networks were built on the data matrices from 1) an LB subset of cases including those with neuropathological diagnoses of control, PD, PDD, and DLB and 2) an AD subset of cases comprising controls and AD. These subset data matrices also included GIS data. Two additional validation networks were also built on the 44 Emory and 103 ROSMAP cases. For each build, network connectivity outlier removal was performed as described [17, 18, 20]. The WGCNA::blockwiseModules() function was used to generate each network. The UPenn and Emory LB networks were built using the following WGCNA settings: soft threshold power = 11.0, deepSplit = 2, minimum module size = 25, merge cut height = 0.07, mean topological overlap matrix (TOM) denominator, a signed network with partitioning about medioids (PAM) respecting the dendrogram, and a reassignment threshold of *p* < 0.05 with clustering completed within a single block. Soft threshold power was adjusted in the UPenn AD and ROSMAP LB networks to 12.0 and 7.5, respectively. As previously described [17], this function generates a correlation matrix across all proteins within each network and subsequently clusters proteins hierarchically into modules based on protein expression pattern similarity across samples. Module eigenproteins are also generated, each representing the first principal component or weighted expression profile of its respective module. The signedkME function of WGCNA then allowed us to determine the bicor correlation between each individual protein and each module eigenprotein. This measure of module membership is defined as kME and was ultimately utilized to determine hub status [51]. To enforce a kME table with no aberrant module assignments, a post-hoc clean-up procedure with iterative protein reassignments were performed as previously described [17, 20].

### Gene Ontology and Cell Type Marker Enrichment Analyses

Gene ontology (GO) annotations were retrieved from the Bader Lab’s monthly updated. GMT formatted ontology lists as previously described [96]. A Fisher’s exact test for enrichment was performed into each module’s protein membership using an in-house script. Molecular function, biological process, and cellular compartment assignments for each module were determined using the highest ranked GO terms associated with each module. Cell type enrichment was also investigated as previously described [17, 20] by analyzing module overlap with RNA sequencing (RNA-seq) and proteomic reference lists of cell type-specific markers [97, 98]. Fisher’s exact tests (FET) were performed to measure the extent of cell type enrichment in each module and were corrected by the Benjamini-Hochberg procedure.

### GWAS Module Association

The enrichment of UPenn LB modules with GWAS targets was performed using MAGMA (version 1.08b) [45] and disease-specific single nucleotide polymorphism (SNP) summary statistics, as previously described [17, 20]. PD summary statistics were derived from http://www.pdgene.org, while AD summary statistics were obtained from Kunkle et al [99]. These lists were filtered for those genes with disease association values of *p*<0.05 prior to enrichment analyses.

### Betweenness Centrality Calculations

To determine the bottleneck status of SNCA and other proteins in its assigned module (M19) in the UPenn LB network, we calculated the betweenness centrality (*g*) of each of these proteins relative to related presynaptic modules, including M7, M17, and M26. Essentially, these betweenness values represent the number of shortest paths passing through a certain protein or node, and those nodes with high betweenness are responsible for the flow of information in that portion of the network [49, 50, 52]. These betweenness measures were calculated in NetworkX (v3.1) in Python (v3.11.5) using the topological overlap matrix (TOM) plot generated by WGCNA upon building the UPenn LB network. A subgraph was constructed using the protein members of M19, M7, M17, and M26. Edges with weights less than the mean edge weight were removed to focus on proteins with the most similar expression patterns. The betweenness centrality was calculated for the resultant graph with protein members of M19 as source nodes and the remaining proteins in M7, M17, and M26 as target nodes. After betweenness centrality was calculated, proteins in M19 were ranked.

### Network Module Preservation

The WGCNA::modulePreservation() function was used to assess network module preservation across networks, as previously described [17, 18, 20]. This function generated a Z_summary_ composite score for each module, using one designated network as the template for each pairwise network comparison. We also assessed module preservation using synthetic eigenproteins as previously published [17, 18, 20]. Briefly, using one network as a template, synthetic modules were assembled in the comparison network comprising the top 20^th^ percentile of proteins by kME. The WGCNA::moduleEigengenes() function was then used to calculate the weighted eigenproteins of these synthetic modules, representing the variance of all synthetic module members across disease cohorts.

### Other Statistics

Statistical analyses were performed in R (version 3.5.2). Correlations were performed using the biweight midcorrelation function as implemented in the WGCNA R package. Comparisons between two groups were performed by *t* test. Comparisons among three or more groups were performed with one-way ANOVA with Tukey or Bonferroni correction for multiple pairwise comparisons of significance. *P* values were adjusted for multiple comparisons by false discovery rate (FDR) correction where indicated. Box plots represent the median and 25th and 75th percentiles, while data points up to 1.5 times the interquartile range from each box hinge define the extent of error bar whiskers. Data points outside this range were identified as outliers.

## Supplementary Materials

**Figure S1. Module overlap between UPenn and Emory co-expression networks.** A hypergeometric Fisher’s exact test was used to determine which modules shared significant overlap of protein members between the UPenn and Emory networks. The 33 modules in the UPenn network (x-axis) were aligned to the 39 modules in the Emory network (y-axis). Numbers indicate the -log10 *p*-value with red shading indicating the degree of significance of overlap (*, *p*<0.05; **, *p*<0.01; ***, *p*<0.001). The green box indicates those UPenn modules that overlapped most strongly with E-24, which stood out in the Emory network given its association with synaptic ontologies, robust elevations in LBD, and selectively positive correlation to LB deposition relative to amyloid and tau. Its overlapping UPenn modules included M17 and M26, which were also SNCA-associated synaptic modules with strong increases in LBD.

**Figure S2. LBD-associated network alterations are replicated in a ROSMAP tissue cohort. (A)** Module preservation analysis of UPenn network into the ROSMAP network. Modules with a Z_summary_ score of greater than or equal to 1.96 (*q*=0.05, blue dotted line) were considered preserved, while modules with Z_summary_ scores of greater than or equal to 10 (*q*=1.0E-23, red dotted line) were considered highly preserved. **(B)** Select UPenn LBD network module eigenproteins associated with their corresponding synthetic eigenproteins in the ROSMAP network. The ROSMAP synthetic eigenproteins reflected the weighted module abundance of the top 20% of proteins by kME comprising each LBD module. ANOVA *p* values are provided for each eigenprotein plot. Box plots represent the median and 25th and 75th percentiles, while data points up to 1.5 times the interquartile range from the box hinge define the extent of error bar whiskers. Abbreviations: CTL, control; PD, Parkinson’s disease; PDD, Parkinson’s disease dementia; DLB, Dementia with Lewy bodies; AsymLB, Asymptomatic Lewy body pathology; LBD, Lewy body dementia.

Table S1: Summary of UPenn cohort demographics and clinicopathological traits

Table S2: Differential protein abundance across UPenn LBD cases

Table S3: UPenn LBD network module assignments

Table S4: Gene ontology z-scores for UPenn LBD network modules

Table S5: PD GWAS MAGMA in UPenn LBD network

Table S6: AD GWAS MAGMA in UPenn LBD network

Table S7: Bottleneck rankings for M19 proteins among UPenn presynaptic modules

Table S8: UPenn Protein correlations to synuclein (SNCA) abundance

Table S9: Summary of Emory cohort demographics and clinicopathological traits

Table S10: Emory LBD network module assignments

Table S11: Gene ontology z-scores for Emory LBD network modules

Table S12: Summary of ROSMAP cohort demographics and clinicopathological traits

Table S13: Differential protein abundance across high- and low-amyloid UPenn LBD cases

## Acknowledgments

We are grateful to those who agreed to donate their brains for research and who participated in the described observational studies. This study was supported by the following National Institutes of Health funding mechanisms: K23NS119964-01 (L.H.), R21NS123882-01 (L.H.), U01AG061357 (N.T.S.) and 1U01NS128433-01 (N.T.S. and A.I.L.). Procurement of UPenn tissues was supported by the following grants: P30AG072979, P01AG066597, U19AG062418 (E.B.L.).

## Author Contributions

Conceptualization, A.S., A.I.L., N.T.S., and L.H.; Methodology, A.S., E.B.D., D.M.D., N.T.S., and L.H.; Investigation, D.M.D., L.Y., and L.H.; Formal Analysis, A.S., E.B.D., O.U., E.K.C., and L.H.; Writing – Original Draft, A.S. and L.H..; Writing – Review & Editing, A.S., E.B.D., E.K.C., D.M.D., J.J.L., A.I.L., N.T.S., and L.H.; Funding Acquisition, A.I.L., N.T.S., and L.H.; Resources, M.G., A.C.P., E.B.L., D.A.B.; Supervision, A.I.L., N.T.S., and L.H.

## Data and Code Availability

Raw mass spectrometry data from the ROSMAP dorsolateral prefrontal cortex tissues can be found at https://www.synapse.org/#!Synapse:syn53177242. Pre- and post-processed protein expression data and case traits related to this manuscript are available at https://www.synapse.org/#!Synapse:syn53177242. The algorithm used for batch correction is fully documented and available as an R function, which can be downloaded from https://github.com/edammer/TAMPOR. Other algorithms implemented in R for cell type marker enrichment (https://github.com/edammer/CellTypeFET), GO annotation (https://github.com/edammer/GOparallel), MAGMA gene level risk enrichment (https://github.com/edammer/MAGMA.SPA), and bicor or standard volcano statistic calculation and plotting (https://github.com/edammer/parANOVA) are also available online. The results published here are in whole or in part based on data obtained from the AMP-AD Knowledge Portal (https://adknowledgeportal.synapse.org). The AMP-AD Knowledge Portal is a platform for accessing data, analyses, and tools generated by the AMP-AD Target Discovery Program and other programs supported by the National Institute on Aging to enable open-science practices and accelerate translational learning. The data, analyses, and tools are shared early in the research cycle without a publication embargo on secondary use. Data are available for general research use according to the following requirements for data access and data attribution (https://adknowledgeportal.synapse.org/#/DataAccess/Instructions).

