## Supplementary figures and images for "Network Proteomics of the Lewy Body Dementia Brain Reveals Presynaptic Signatures Distinct from Alzheimer’s Disease"

### Supplemental Figures

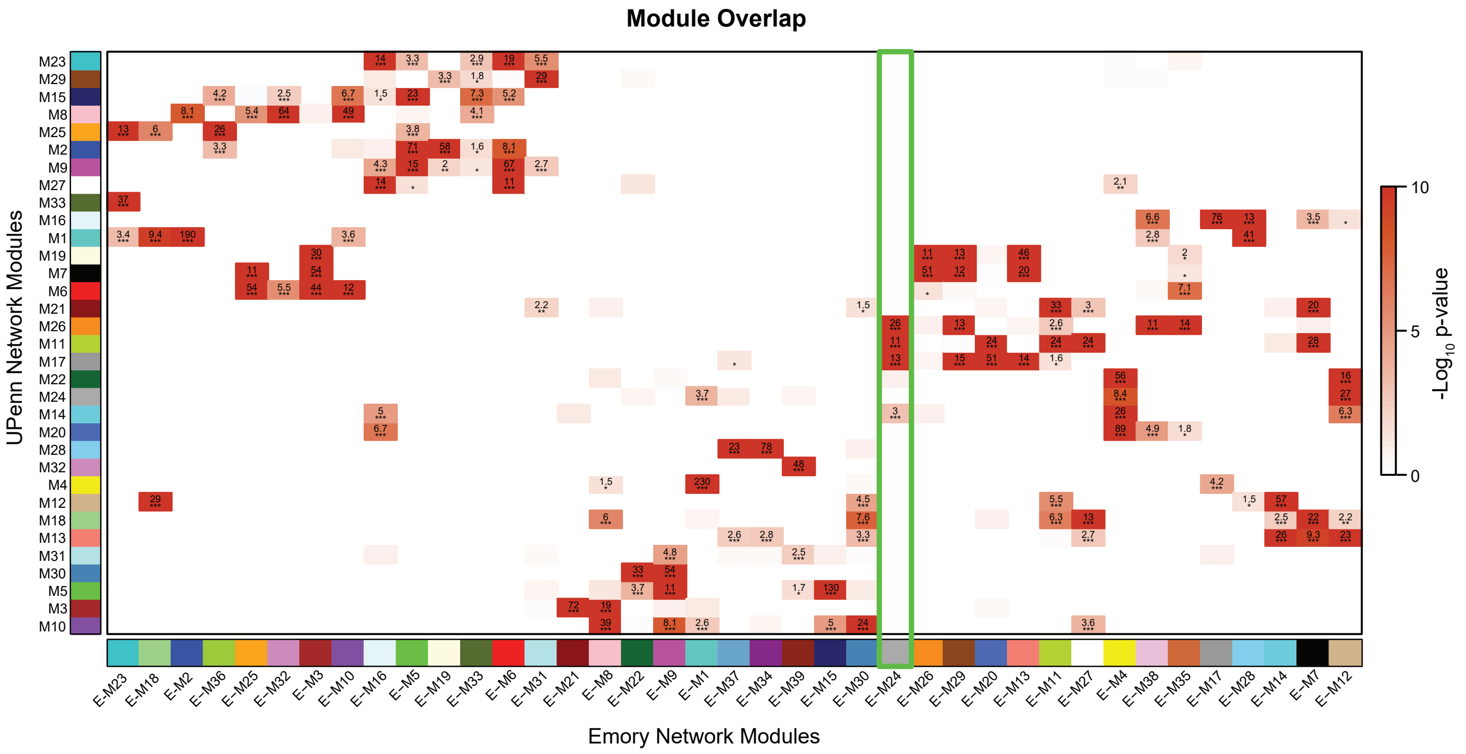

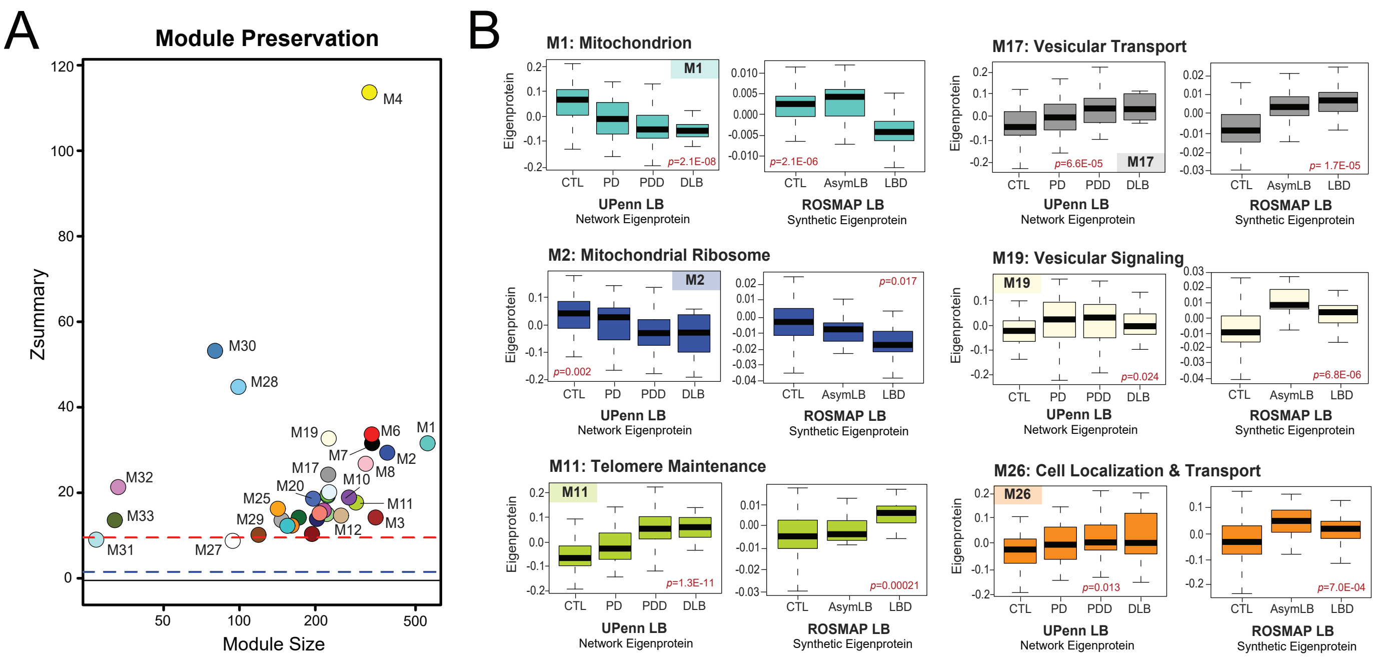
